# Mouse brain organoids model *in vivo* neurodevelopment and function and capture differences to human

**DOI:** 10.1101/2024.12.21.629881

**Authors:** Daniel J Lloyd-Davies Sánchez, Feline W Lindhout, Alexander J Anderson, Laura Pellegrini, Madeline A Lancaster

## Abstract

In the last decade since their emergence, brain organoids have offered an increasingly popular and powerful model for the study of early development and disease in humans. These 3D stem cell-derived models exist in a newer space at the intersection of *in vivo* and 2D *in vitro* models. Functional benchmarking has so far remained largely uncharacterised however, leaving the extent to which these models may accurately portray *in vivo* processes still yet to be fully realised. Here we present a standardised unguided protocol to generate brain organoids from mice, the most commonly-used *in vivo* mammalian model; and in parallel establish a guided protocol for generating region-specific choroid plexus mouse organoids. Both unguided and guided mouse organoids progress through neurodevelopmental stages with an *in vivo*-like tempo and recapitulate species-specific characteristics of neural and choroid plexus development, respectively. Neuroepithelial cells generate neural progenitors that give rise to different neural subtypes including deep-layer neurons, upper-layer neurons, and glial cells. We further adapted protocols to prolong mouse cerebral organoid (CO) cultures as slices at the air-liquid interface (ALI), enhancing accessibility for long-term studies and functional investigations. In mature mouse ALI-COs, we observed mature glia, as well as synaptic structures and long-range axon tracts projecting to distant regions, suggesting an establishment and maturation of neural circuitry. Indeed, functional analyses with high-density multi-electrode arrays (HD-MEAs) indicate comparable activity to *ex vivo* organotypic mouse brain slices. Having established protocols for both region-specific and unpatterned mouse brain organoids, we demonstrate that their neurodevelopmental trajectories, and resultant mature features, closely mimic the *in vivo* models to which they are benchmarked across multiple biochemical, morphological, and functional read-outs. We propose that mouse brain organoids are a valuable model for functional studies, and provide insight into how closely brain organoids of other species, such as human, may recapitulate their own respective *in vivo* development.

## Introduction

Biochemical and medical research utilises a plethora of *in vitro* and *in vivo* model systems to study biology from the molecular to the organismal scale. However, technical differences between *in vitro* and *in vivo* systems, and the biological differences between humans and model species frequently frustrate the pipeline of fundamental research to its application in pharmaceutical industry and technology ^1^. The study of human brain development is especially difficult to model due to the marked differences in size and complexity of primate brains compared to other mammals ^2,3^, plus the limited availability and intervention possible with human-derived tissue, among other obstacles ^4^.

In the last decade, the use of stem cell-derived 3-dimensional neural organoid models has accelerated *in vitro* research of human neurodevelopment ^5^. A variety of protocols for organoid generation allows for the study of endogenous developmental programs ^6^, or the production of specific neural cell types and brain regions ^7–11^ from human cell lines. Moreover, the use of gene-editing and patient-derived induced pluripotent stem cells (iPSCs) has enabled the modelling of diseases whose phenotypes are poorly or only partially replicated in animal models, such as microcephaly ^6,12^, Alzheimer’s disease ^13,14^ and neuronal heterotopia ^15^. Additionally, neural organoids afford greater experimental intervention and control at a lower cost than *in vivo* vertebrate studies ^16,17^. To ensure their applicability as a model, benchmarking studies have sought to characterise the fidelity of organoid culture to *in vivo* development. Many studies have compared molecular signatures to establish that diverse cell identities, their developmental dynamics, and cytoarchitecture are reproduced in organoids, with minor differences ^18^, via epigenomics ^19–22^, transcriptomics ^23–26^, proteomics ^27^, and immunostaining of cell type specific markers. Whereas a limited number have shown electrophysiological function in organoids via calcium imaging ^28,29^, patch-clamping ^30,31^ and multi-electrode array electrophysiology ^11,25,32^, the relevant stage-matched human comparison is currently limited to non-invasive electroencephalography (EEG) of preterm newborns ^33,34^ or *ex vivo* cortex slice culture ^33,35^ which is scarce. Thus, benchmarking efforts are limited when comparing cell maturity, behaviour, and network activity due to the same challenges that motivate the use of *in vitro* models to study human fetal brain development. To achieve robust functional benchmarking of neural organoids, alternative approaches integrating *in vivo* and *in vitro* comparisons are required, allowing for more detailed analysis using diverse methodologies.

The mouse (*Mus musculus*) is the most widely used mammalian *in vivo* model, underpinning seminal neurodevelopmental findings and near ubiquitous in early-stage pharmaceutical trials. As such, there is a wealth of *in vivo* data, from molecular signatures ^36,37^ to neurophysiology ^38–40^, and established protocols to derive such information. This enables effective benchmarking of new and more complex *in vitro* systems, such as neural organoids, to discern what artefacts, insufficiencies or deficiencies are present in these models compared to the *in vivo* biology they model. Furthermore, the combination of human and mouse *in vitro* organoid models may bridge between *in vivo* mouse experiments and human clinical trials which so often result in costly failure, particularly for neurological treatments ^41,42^. As such, there has been a surge of interest in establishing methods to generate mouse brain organoids, with protocols now available for mouse cerebellar organoids ^7^, mouse hippocampus organoids ^43,44^, as well as mouse epiblast stem cell-derived forebrain organoids ^45,46^.

In this study, we report mouse cerebral organoids generated from murine embryonic stem cells using the same reagents and unguided differentiation approach as employed in generating human and non-human primate organoids. We also adapted a patterned approach previously used to generate human choroid plexus organoids, to develop region-specific mouse choroid plexus organoids. We then focus on cerebral organoids and show that mouse organoids enter neurogenesis sooner than human organoids and generate diverse neuronal and glial cell types with a faster developmental tempo. We detail their later-stage development in air-liquid interface culture and observe the maturation of synaptic structures, the formation of long-range axon tract bundles, and activity of functional neuronal units which emulate those of *ex vivo* mouse brain slices. This work demonstrates that unguided organoids recapitulate the tempo and hallmarks of *in vivo* neuronal maturation and physiology in a complex tissue context *in vitro*. The rapid development of mouse organoids can facilitate high-throughput studies of later neurodevelopmental stages with promising fidelity. They may also hasten the development of new organoid protocols or analyses and provide an outgroup for multispecies evolutionary studies in organoids. Overall, mouse cerebral and choroid plexus organoids complement findings *in vivo* from mice, and human organoid models, improving the generalisation and translation potential of neurodevelopmental research to human biology and health.

## Results

### Generation of unguided mouse cerebral organoids

We sought to generate mouse cerebral organoids to discern whether discordant findings from human organoids and *in vivo* mouse models can be attributed to species-specific differences or artefacts of *in vitro* culture. We successfully generated unguided mouse cerebral organoids by modifying the timing of our previously established protocol for human cerebral organoid culture while keeping all other aspects, including media formulations, identical (**Figure 1A**). In prior studies, we had observed that human and non-human ape organoids mature at species-specific rates reflective of differences in the timing of their *in vivo* development ^47^. Consistent with the rapid neural development of mice *in vivo* ^48^, mouse cerebral organoids established large rosette structures within 7 days (**Figure 1B**). This was notably faster than current human organoid protocols under the same conditions. By monitoring size, morphology, the presence of non-neural lineages, and attainment of differentiated cell types, we empirically determined protocol timings for the robust generation of mouse forebrain tissue.

**Figure 1:**
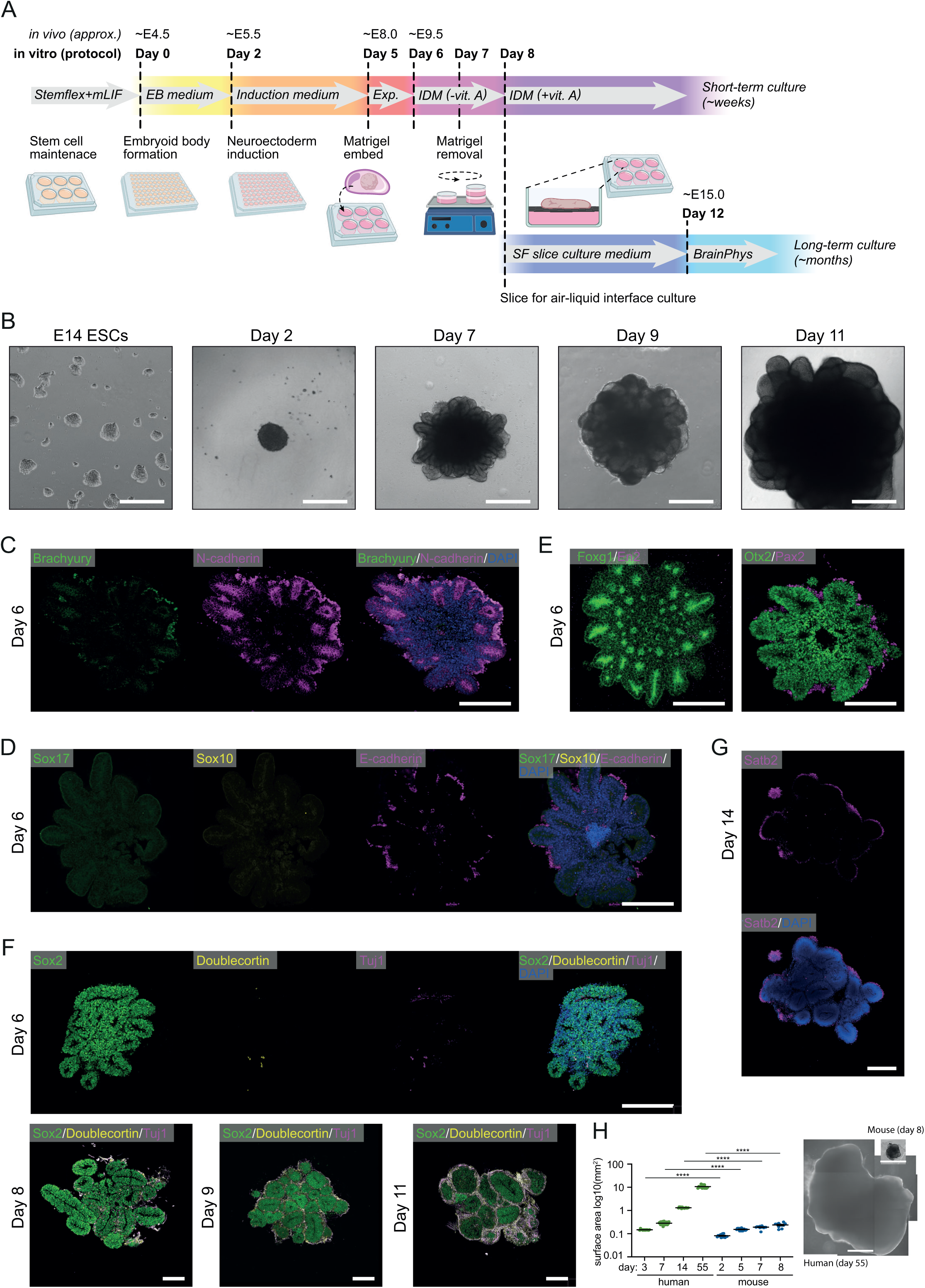
Generation of mouse unguided cerebral organoids with forebrain identity. A) Schematic of the mouse organoid culture protocol as detailed in the materials & methods. Media formulations, culture dishes and interventions are indicated for each timepoint. Between day 8 and 12 of the protocol, organoids may be sliced and cultured at the air-liquid interface for long-term maintenance, or maintained as unsliced organoids on an orbital shaker. Stages of early organoid development are matched to mouse developmental milestones derived from *in vivo* studies (E = embryonic day). mLIF, mouse leukemia inhibitory factor; EB, embryoid body; Exp., Expansion medium; IDM, Improved Differentiation Medium; vit. A, vitamin A; SF, serum-free. B) Brightfield images of mouse organoid development. E14 embryonic stem cells (ESCs) are harvested to form embryoid bodies (Day 2) and form neuroepithelial buds once embedded in Matrigel droplets (Day 7). Subsequently, Matrigel is completely removed (Day 9) and the organoid grows rapidly in size (Day 11). Scale bar = 500 μm. Representative image of n = 20 organoids in two independent batches. C) Immunohistochemical staining for germ layer lineage markers in day 6 mouse organoids. Brachyury indicative of neuromesoderm, and N-Cadherin (magenta) indicative of neuroepithelium. Nuclei stained with DAPI (blue). Scale bar = 200 μm. Representative image of n = 4 organoids in two independent batches. D) Immunohistochemical staining for germ layer lineage markers in day 6 mouse organoids. Sox17 (green) indicative of endoderm, Sox10 (yellow) indicative of neural crest, and E-Cadherin (magenta) indicative of non-neural epithelium. Nuclei stained with DAPI (blue). Scale bar = 200 μm. Representative image of n = 3 organoids. E) Immunohistochemical staining for markers of brain regionalisation in day 6 mouse organoids. Left: Foxg1 (green) indicative of telencephalon, and En2 (magenta) indicative of cerebellum. Right: Otx2 (green) indicative of forebrain and midbrain, and Pax2 (magenta) indicative of midbrain and anterior hindbrain. Nuclei stained with DAPI (blue). Scale bar = 200 μm. Representative image of n = 3 organoids. F) Immunohistochemical staining for cell identity markers in day 6, day 8, day 9 and day 11 mouse organoids. Sox2 (green) indicative of neuroepithelia and radial glia, doublecortin (yellow) and Tuj1 (magenta) indicative of neurons. Nuclei stained with DAPI (blue). Scale bars = 200 μm. Representative image of n = 3-4 organoids per timepoint. G) Immunohistochemical staining for a cell identity marker in day 14 mouse organoids. Satb2 (magenta) indicative of upper-layer neurons. Nuclei stained with DAPI (blue). Scale bar = 500 μm. Representative image of n = 4 organoids. H) Quantifications of human and mouse brain organoid size across different steps of the protocol. Human day 3: n = 14 organoids, human day 7: n = 30, human day 14: n = 15, human day 55: n = 14, mouse day 2: n = 15, mouse day 5: n = 15, mouse day 7: n = 15, mouse day 8: n = 11. All data are shown by mean ± SEM; \*\*\*\**P* < 0.0001 as determined by unpaired *t* tests with Bonferroni correction. Representative images of human day 55 and mouse day 8 organoids to scale. Scale bar = 1mm

On day 6 of our mouse-specific protocol (**Figure 1A**), rosette structures showed N-cadherin expression at their apical surface, a marker of neuroepithelium (**Figure 1C**). Moreover, the organoids were negative for the expression of Brachyury, Sox17, and Sox10 which indicate mesoderm, endoderm, and neural crest, respectively (**Figure 1C-D**). Immunostaining for E-cadherin, a marker of non-neural ectoderm, was positive for a small number of cells toward the centre of a few organoids. Overall, the cells of the developing organoids were highly restricted to a neural ectoderm lineage as desired. Day 6 organoids were also positive for forebrain markers Foxg1 and Otx2 but negative for the hindbrain marker Pax2, and cerebellar marker En2 by immunostaining (**Figure 1E**) (other than for weak signal seen in extra-organoid debris and Matrigel autofluorescence which is lost at later timepoints after Matrigel removal). By day 6 of our protocol (**Figure 1A**), we have robustly generated early forebrain cell identities to the exclusion of other germ layers and brain regions.

The onset of neurogenesis was first observed in mouse cerebral organoids on day 6. Neural rosettes predominantly consisted of Sox2-positive radial glia/neuroepithelia, however a small number of immature neurons were identified by Doublecortin and Tuj1 immunostaining (**Figure 1F**). The number of Doublecortin and Tuj1 positive neurons increased dramatically between day 6 and day 11, whilst a distinct progenitor zone of Sox2-positive cells was maintained (**Figure 1F**). By day 14, we observed Satb2 expression indicating the presence of mature neuronal subtypes and the onset of upper-layer neurogenesis in the patterning of a cortical plate (**Figure 1G**). From the timing and expression patterns of these early-stage mouse organoids we could estimate the corresponding *in vivo* developmental time as reported in the literature ^49–52^ (**Figure 1A**). Further reflective of species-specific differences seen *in vivo*, we observed that the faster-developing mouse cerebral organoids grew smaller than their human organoid counterparts at comparatively benchmarked stages (**Figure 1H**).

### Generation of region-specific, patterned choroid plexus mouse organoids

Functional benchmarking of brain organoids across species requires complementary models that allow direct comparison with *in vivo* data. To generate brain region-specific mouse organoids we focused on the choroid plexus (ChP): a secretory tissue located in each ventricle of the brain. The ChP is composed of a single layer of epithelial cells surrounding a core of connective tissue, stromal and immune cells ^53–55^. This tissue is responsible for the production of cerebrospinal fluid (CSF) and formation of the blood-CSF-barrier (**Figure 2A**). Previously, we generated human ChP organoids that secrete CSF and form a barrier ^8^ using patterning factors that drive ChP differentiation ^56,57^. To generate mouse ChP organoids, we adapted the same approach and patterning factors used for human ChP organoids.

**Figure 2:**
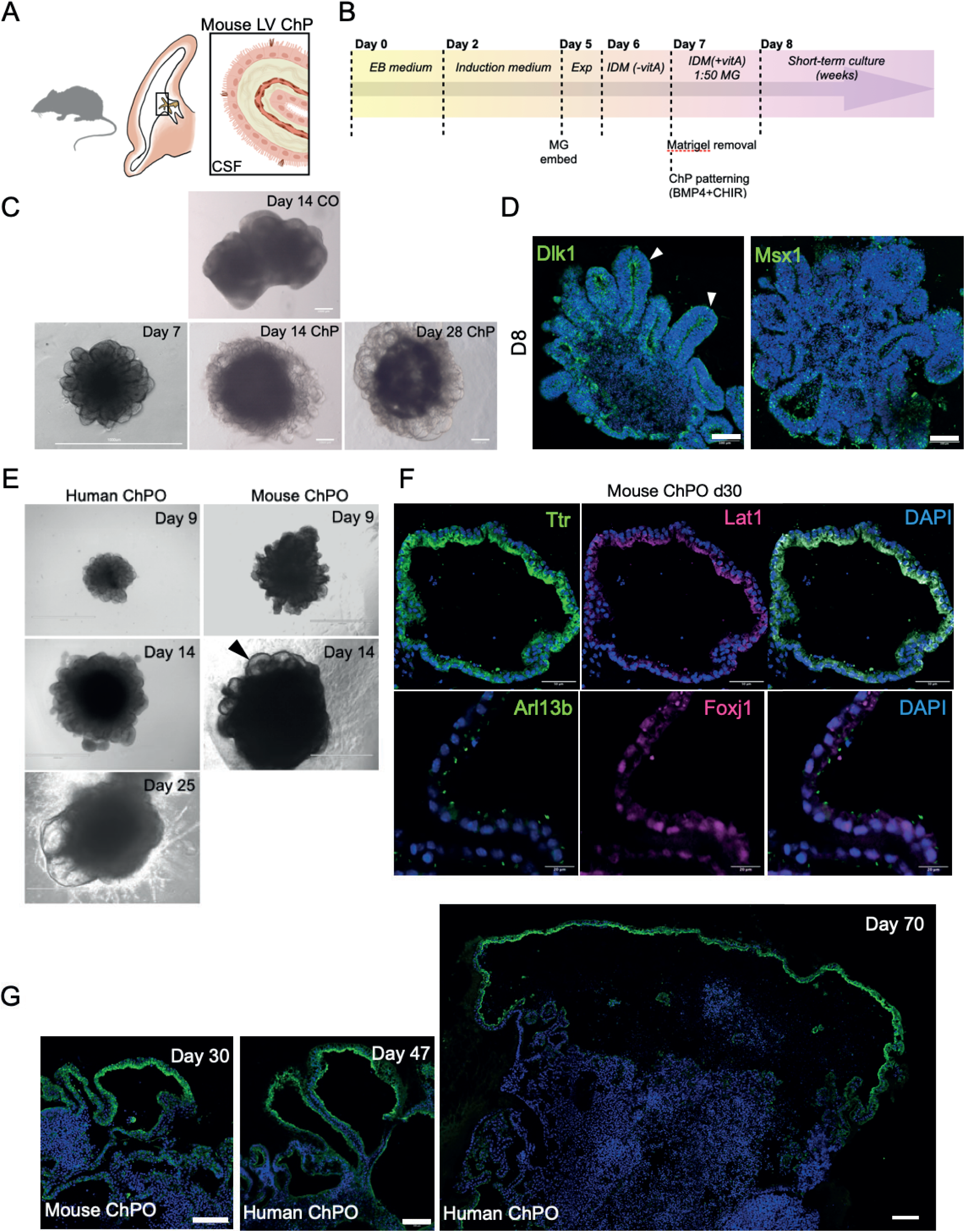
Generation of mouse region-specific choroid plexus organoids with multiciliated cells. A) Schematic of mouse lateral ventricle (LV) choroid plexus with ciliated cells and producing cerebrospinal fluid (CSF). B) Protocol timeline for the mouse choroid plexus organoid culture as detailed in the materials & methods. Media formulations, culture dishes and interventions are indicated for each timepoint. Choroid plexus patterning molecules BMP4 and CHIR are applied on day 7. The maturation media from day 7 is IDM supplemented with vitamin A and dissolved matrigel (MG). mLIF, mouse leukemia inhibitory factor; EB, embryoid body; Exp., Expansion medium; IDM, Improved Differentiation Medium; vit. A, vitamin A; SF, serum-free. C) Brightfield images of day 14 cortical (CO) and choroid plexus (ChP) mouse organoids from the same batch. Scale bar = 1000 μm. Representative images of n = 4-5 organoids in three independent batches. D) Immunohistochemical staining for cell identity markers of day 9 ChP mouse organoids. Dlk1 (green) indicative of cortical neuroepithelium and Msx1 (green) indicative of early ChP and cortical hem. Scale bar = 100 μm. Representative images of n = 2-3 organoids in two independent batches. E) Immunohistochemical staining for cell identity markers of day 30 ChP mouse organoids. Ttr (green) indicates ChP specification, Lat1 (magenta) stains for an aminoacid transporter expressed in the ChP. Scale bar = 50 μm. The panel below shows ciliated cells stained with FOXJ1 (magenta) and Arl13b (green). Scale bar = 20 μm. Representative images of n = 2-3 organoids in two independent batches. F) Immunohistochemical staining for cell identity markers of day 30 ChP mouse organoids. Ttr (green) indicates ChP specification, Lat1 (magenta) stains for an aminoacid transporter expressed in the ChP. Scale bar = 50 μm. The panel below shows ciliated cells stained with Foxj1 (magenta) and Arl13b (green). Nuclei stained with DAPI (blue). Scale bar = 20 μm. Representative images of n = 2-3 organoids in two independent batches. G) Immunohistochemical staining for Ttr (green) of day 30 mouse ChP organoids in comparison with human H1 ChP organoids at day 47 and day 70. Nuclei stained with DAPI (blue). Scale bar = 100 μm.

The protocol to generate mouse ChP organoids follows the same initial steps as the unguided cerebral organoid protocol. After neuroepithelium generation and expansion at day 7, EBs were patterned with BMP4 and CHIR to promote ChP differentiation (**Figure 2B**). After one day of treatment, it was possible to observe the elongated shape of the neuroepithelial buds, a hallmark of ChP morphogenesis, and development of a polarised epithelial monolayer expressing early ChP markers such as Msx1 (**Figure 2C,D**). After 14 days, mouse organoids treated with BMP4 and CHIR appeared completely enriched in ChP epithelium compared to untreated, unguided cerebral organoids (**Figure 2C**).

In mice, the ChP begins to develop around embryonic day 10 (E10) with the emergence of the telencephalic ChP, followed by hindbrain and third ventricle ChP by E12 ^54,55^. By E14, these structures are morphologically distinct and actively produce CSF. In humans, the ChP appears around gestational week 6-7, with a similar sequence of formation in the lateral, third and fourth ventricles. By the end of the first trimester (12 weeks), the human ChP is well-formed and contributing to CSF secretion ^55^. Compared to mice, and similarly to other brain regions, the human ChP development is also prolonged and leads to a more complex and expanded structure. Consistently, mouse ChP organoids appear to already produce the characteristic CSF-producing compartments around day 14, whereas human ChP organoids start producing CSF only around day 20-25 (**Figure 2E**).

To validate the presence of specific ChP markers we performed histological analysis of day 30 mouse ChP organoids, which revealed expression of the ChP marker Ttr and amino acid transporter Lat1 (**Figure 2F**). These organoids also developed distinct cellular populations, including multiciliated epithelial cells expressing Foxj1, a transcription factor required for formation of multiciliated epithelium, and with cilia positive for Arl13b (**Figure 2F**). We also observed that mouse ChP organoids reached a maximum size around day 30 and appeared much smaller, forming less expanded fluid-filled cysts, compared to the human ChP organoids, reflecting species-specific differences in ChP structure (**Figure 2G**). Importantly, this provides a proof-of-principle model for generating region-specific brain organoids that align with *in vivo* developmental timelines, offering a valuable tool for future functional studies.

### Neuronal maturation in unpatterned mouse ALI-COs

To investigate maturation stages of mouse cerebral organoids, we adopted a protocol allowing for their long-term maintenance by preparing 300 µm organoid sections, thus several cell layers thick, that were kept at the air-liquid interface. This method is classically employed for prolonged *ex vivo* mouse brain cultures, which are similarly maintained as organotypic slice cultures at the air-liquid interface, enabling not only their long-term maintenance but also the accessibility of various brain structures for live imaging and electrophysiological analysis. More recently, this protocol was adapted and modified for long-term maintenance of human brain organoids, enabling prolonged and improved survival for several months ^32^. We extended this technique to mouse brain organoids to model later maturation stages. Slice cultures prepared from mouse brain organoids between days 8 and 13 were maintained for weeks to months, with the possibility for further extending durations if required (**Figure 3A**).

**Figure 3:**
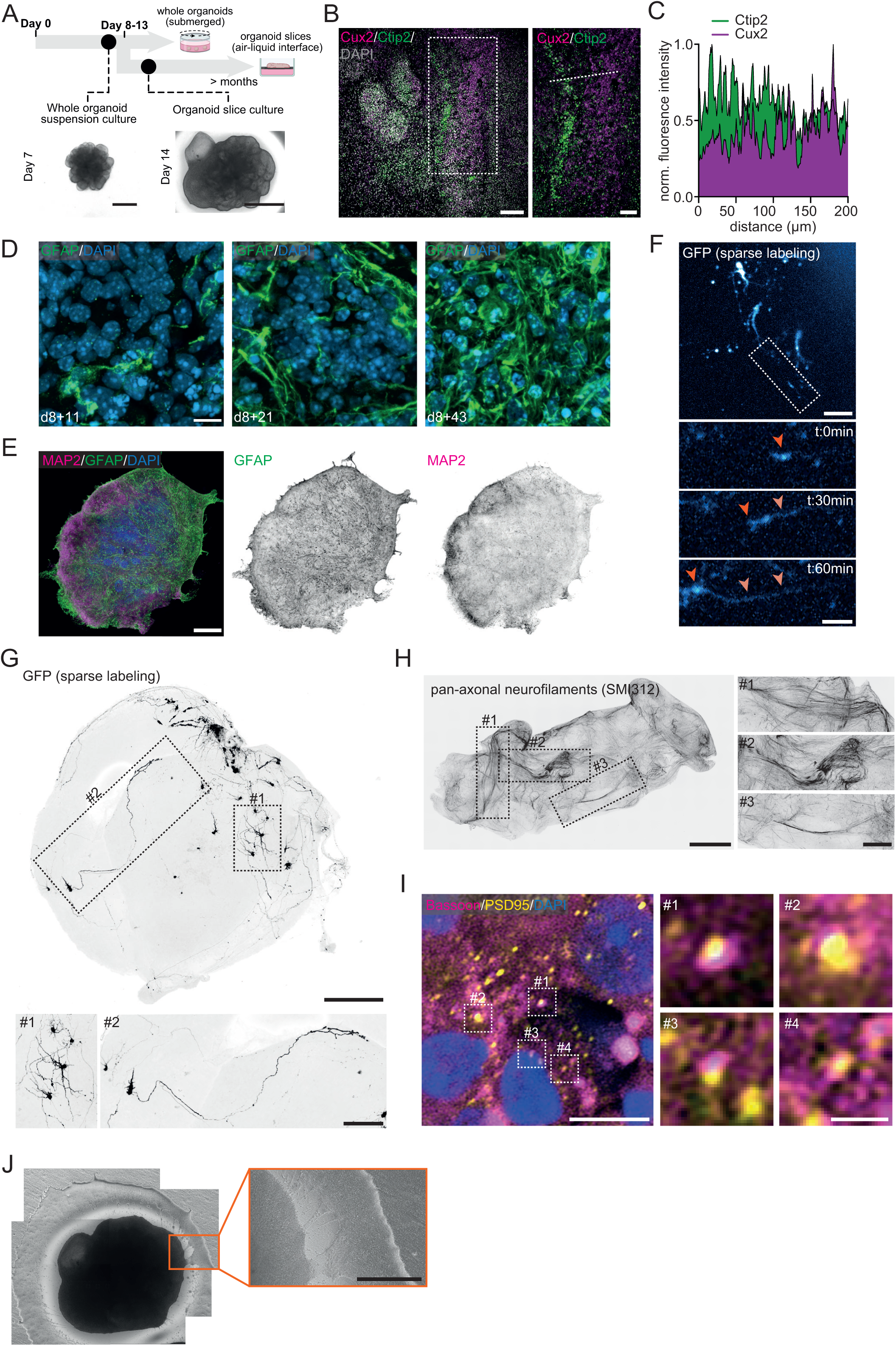
Long-term mouse brain organoid slice cultures display hallmarks of neuronal maturation. A) Top: schematic illustration of the generation of air-liquid interface slice cultures from day 8-12 mouse brain organoids. Bottom: brightfield images of a brain organoid (unsliced) at day 7 (scale bar = 250 μm) and a brain organoid slice culture at the air-liquid interface at day 12+2 (scale bar = 1000 μm). B) Cryosectioned mouse brain organoid slice culture at day 12+46 immunostained for Cux2, marking deep layer neurons, Ctip2, marking upper layer neurons, and DAPI, marking nuclei. Zoom displays a region within the organoid displaying spatial segregation of deep and upper layer neurons, as found in various areas of the brain organoid slice cultures. Scale bars = 100 μm (full size) and 50 μm (zoom). Representative image of n=3 organoids in 2 independent batches. C) Normalised fluorescent intensity plot of indicated line (dotted) in (B). D) Cryosectioned mouse brain organoid slice cultures at day 8+11, 8+21 and 8+43, immunostained for GFAP, marking glia cells, and DAPI, marking nuclei. Scale bar = 10 μm. E) Whole-mount mouse brain organoid slice culture at day 8+46 immunostained for MAP2, marking neurons, GFAP, marking glia cells, and DAPI, marking nuclei. Scale bar = 500 μm. Representative image of n=3 organoids from 1 batch. F) Timelapse of a mouse brain organoid slice culture at day 8+8 with sparsely GFP-labelled cells using Sendai virus. Zoom shows a growing axon tip, indicated by orange arrowheads. Scale bars: 50 μm (full size) and 20 μm (zoom). Representative timelapse of n=8 organoids in 3 independent batches. G) Whole-mount mouse brain organoid slice culture at day 8+7 with sparsely GFP-labelled cells using Sendai virus. Zooms display examples of individual neurons with mature morphologies, including extended axons and complex dendritic arborizations. Scale bars = 500 μm (full size) and 200 μm (zoom). Representative image of n=27 organoids in 9independent batches. H) Whole-mount mouse brain organoid slice culture at day 8+14 immunostained for pan-axonal neurofilaments (SMI312), marking the total axon population within the brain organoid slice culture. Zooms display examples of self-assembled axon tracts projecting to distinct regions within the organoid. Scale bars = 500 μm (full size) and 200 μm (zoom). Representative image of n=3 organoids from 1 batch. I) Superresolution microscopy image of a cryosectioned mouse brain organoid slice culture at day 8+43 immunostained for Bassoon, marking presynaptic sites, PSD95, marking excitatory postsynaptic sites, and DAPI, marking nuclei. Zooms display examples of co-appearing Bassoon and PSD95 puncta, indicative for excitatory synaptic connections. a region within the organoid displaying spatial segregation of deep and upper layer neurons, as found in various areas of the brain organoid slice cultures. Scale bars = 5 μm (full size) and 1 μm (zoom). Representative image of n=17 ROIs of two organoids in 1 independent batch. J) Brightfield image of brain organoid slice culture at day 8+19. Zoom indicates externally projecting axons from the slice in culture. Scale bar = 400 μm. Representative image of n=3 organoids from 1 batch.

The slice cultures revealed distinct neuronal subtype populations, marked by the presence of Ctip2-positive deep layer neurons and Cux2-positive upper layer neurons, which in various areas of the organoids were organised in spatially distinct regions (**Figure 3B,C**). To assess the progression of gliogenesis, a hallmark of late neurodevelopment, we assessed the appearance of GFAP-positive cells with a stellate morphology typical for astrocytes over time. In organoids sliced at day 8, glial cells first appeared 11 days after slicing (8+11) and their abundance progressively increased in subsequent weeks (**Figure 3D,E**). At these later stages, GFAP-positive glial cells, like MAP2-positive neurons, were abundantly present across the brain organoid, displaying mature cellular morphologies as indicated by extended processes (**Figure 3F**).

During the process of neuronal maturation, newborn neurons go through a series of substantial morphological changes, including the formation of long axons and arborized dendrites, and assemble specialised subcellular structures, such as synaptic connections, to ultimately establish functional neuronal networks. We next sought to explore the establishment of these mature neuronal morphologies and structures in more detail. Using a live imaging and sparse viral labelling approach, we observed abundant axonal outgrowth in mouse organoids about a week after slicing (day 8+8) (**Figure 3F**), marking one of the first steps of postmitotic neuronal development. Examining individual neuronal morphologies further showed that neurons exhibited axons extending to local and distal regions, and developed complex and arborized dendrites (**Figure 3G**). Around day 8+14, axons formed self-organised bundles projecting to different regions within the brain organoid slice culture (**Figure 3H**), as well as external projections (**Figure 3J**), as also previously reported in human brain organoid slice cultures ^32^. Of note, to successfully obtain these self-organising axon tracts, it was key to obtain complete Matrigel removal at earlier stages of the protocol, thereby avoiding axonal outgrowth into the Matrigel and away from the organoid (**Supplementary** Figure 1A,B).

To determine the subsequent establishment of synaptic connections, we examined the co-appearance of Bassoon, a presynaptic scaffold protein, and PSD95, a postsynaptic scaffold protein of excitatory synapses, using immunostaining. In mouse organoids ∼6 weeks after slicing (day8+43) we observed an abundance of synaptic structures (**Figure 3I**), indicating the presence of neural networks.

### Functional neuronal activity with *in vivo*-like action potentials in unguided mouse ALI-COs

Mouse organoids exhibited various hallmarks of electrical activity with calcium transients evident in maturing neurons (**Figure 4A, Supplementary Video 1**), and c-fos staining, a common proxy for neuronal activity, demonstrating metabolically active neurons (**Figure 4B**). We therefore examined the electrical activity of these cultures using multielectrode arrays. High spatial resolution of high-density multielectrode arrays (HD-MEAs) enabled construction of heat maps of voltage changes during firing activity, across mouse ALI-CO cultures (**Figure 4C**). Single-cell and subcellular spatial resolution of firing activity recordings facilitated computational clustering of detected action potentials across the HD-MEA and thus reconstruction of the electrical footprint of individual neurons (**Figure 4D**). Being able to reconstruct the electrical footprint of a neuron from locations where action potential voltage changes can be detected is a powerful technique as in this manner we can conduct electrical imaging of neurons (**Figure 4D**) in ALI-COs in a non-invasive way without the need for fixation or processing.

**Figure 4:**
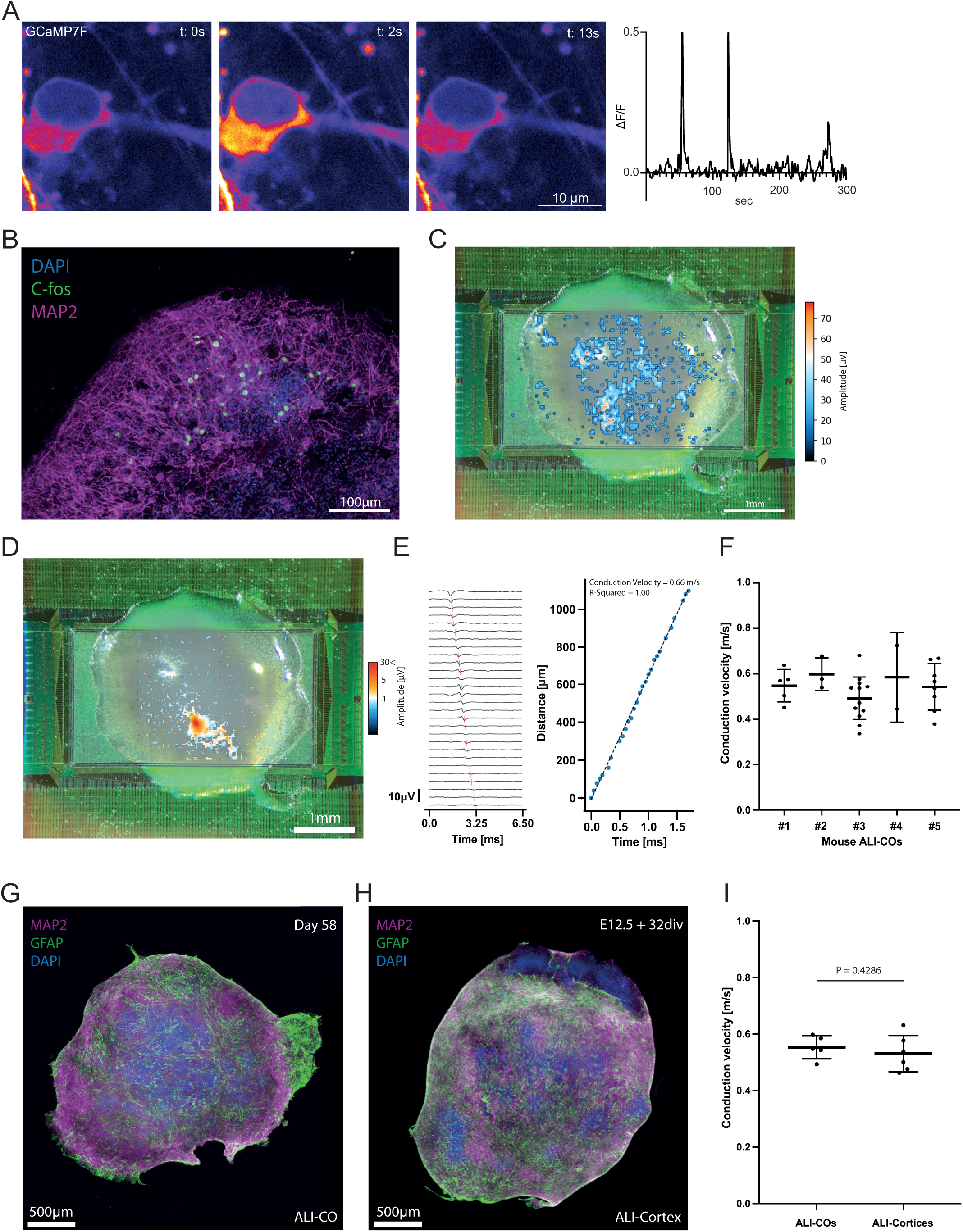
Functional neural circuit establishment in unguided mouse ALI-Cos. A) Left: timelapse of a neuron expressing GCaMP7f in a day 20 mouse brain organoid. Right: ΔF/F0 fluorescent GCaMP7f intensity plot during 5 min time lapse of neuronal soma shown left. B) C-Fos staining reveals metabolically active, possibly electrically active, neurons (MAP2). Scale bar = 100 μm. C) Heatmap of spiking amplitudes for a day 28 mouse ALI-CO recorded on a high-density MEA. Sparse activity scan recording carried out for 5 minutes. Scale bar 1 mm. D) Clustering of action potentials detected on MEA enables reconstitution of electrical footprint of individual neurons with mouse ALI-CO. Scale bar 1 mm. E) Latency of action potential spike at different distances from the axon initial segment of neuron in D. With distance and latency information, conduction velocity of action potentials as they propagate through the axon, can be calculated. F) Mean conduction velocities for individual neurons (the mean of action potentials detected in those neurons during the recording period) are plotted for each mouse ALI-CO recorded. Mean ± SD is shown for the neurons detected within each ALI-CO. G) MAP2 and GFAP stain reveals gross neuronal and glial (respectively) distribution in mouse ALI-CO. Scale bar = 500 mm. H) E12.5 mouse brain slices cultured at the air-liquid interface (ALI-Cortices) exhibit a similar gross appearance to ALI-COs. Scale bar = 500 mm. I) Mean neuronal conduction velocities per ALI-CO shown in F are plotted. Mean ± SD is shown at the level of ALI-COs. E12.5 mouse brain slices cultured at the air-liquid interface for 32 days were recorded in the same way, with comparable data plotted alongside ALI-CO level conduction velocity data. Mann-Whitney test revealed no difference in conduction velocities between ALI-COs and ALI-Cortices (P = 0.4286).

Furthermore, after clustering and assignment of recorded voltage changes to individual neurons, analysis of action potential latency against serial paired data of incremental distances from the point of action potential origin could be performed leading to calculation of a gradient for the conduction velocity of action potentials as they propagate through axons (**Figure 4E**). We found the mean and variation in neuronal conduction velocities were comparable to that of E12.5 mouse brain slices which we had cultured at the air-liquid interface (ALI-Cortices) in the same manner as ALI-COs (**Figure 4F,I**). These data also correlated with published conduction velocities of 0.46m/s for primary E18 rat neurons dissociated and plated on HD-MEAs ^58^, demonstrating that the conduction velocities recorded from organoids are comparable to neurons generated *in vivo* and are not simply artefacts of air-liquid interface culture or the organoid method. Mouse ALI-COs exhibited comparable gross structural appearance (**Figure 4G**) compared to the primary brain tissue cultured in the same way (**Figure 4H**), demonstrating the ability of brain organoid cultures to recapitulate the real developing brain tissue in many ways.

## Discussion

Cerebral organoids and ALI-COs provide a powerful model for neurodevelopment and neuronal maturation due to their 3D nature and recapitulation of a tissue environment. These models offer a more comprehensive platform for modelling neurodevelopmental processes, placing them at the interface of *in vivo* and *in vitro* systems. The presence of different pools of neural progenitors and neural subtypes demonstrates the capacity to generate complex cell diversity, a hallmark of *in vivo* brain development. This spontaneous neurodevelopment ultimately results in mature ALI-COs that exhibit more complex wiring of axon tracts, as shown by the self-organised formation of long-range axon bundles projecting to other regions in the organoid, a phenomenon not observed in 2D cultures. The architecture of axon tracts is further supported by electrophysiological recordings with high spatiotemporal resolution to map axon morphologies, demonstrating functional connectivity. We also show that neurons from unpatterned mouse organoids were able to recapitulate aspects of neuronal physiology, as assessed via comparison to primary mouse embryonic brain slices which had been cultured in the same way. Together, these findings demonstrate the utility of mouse brain organoids as a complementary tool to rodent models for various neuroscience studies and pharmaceutical applications, providing a robust *in vitro* alternative to use across different disciplines and potentially contributing to the reduction of laboratory animals.

Aside from mouse organoids providing a valuable model for bridging the translation of an extensive rodent *in vivo* literature towards a more specific understanding of human brain development, as already discussed; they also provide a valuable reference point and comparison for studies in human and ape organoids particularly through studies of functional networks and neuronal firing. Our data demonstrates mouse organoids and ALI-COs are able to differentiate and generate brain organoid tissue in much the same way as human cerebral organoids ^32,59^ – likely in large part due to the intrinsic nature of differentiation facilitated by an unguided protocol. Indeed, the same media formulations and procedures support neurodevelopmental progression in organoids of each species. Benchmarking these mouse brain organoids against embryonic-derived mouse tissue not only validates their fidelity but also highlights the potential of human brain organoids to accurately model aspects of human brain development and neural circuit establishment. Moreover, the generation of mouse choroid plexus organoids using a guided, region-specific protocol extends the utility of mouse organoid models by offering a complementary system for studying CSF production and early development of the blood-CSF-barrier. These organoids replicate key structural features and functional features of the choroid plexus and align with *in vivo* developmental timings, thereby providing an important reference to cross-species comparison and further bridging the gap between *in vivo* mouse models and human organoids.

Together, these findings highlight the versatility of mouse cerebral and choroid plexus organoids as a tool to model neurodevelopment, functional neuronal networks, and brain-region specific processes. Their ability to complement human brain organoid models and serve as a benchmark for *in vivo* rodent studies positions them as critical models for understanding neurodevelopmental biology while facilitating translational research and reducing reliance on animal models.

## Acknowledgements

The authors would like to thank members of the Lancaster lab for helpful feedback and discussions. We also thank the Light Microscopy facility of the MRC Laboratory of Molecular Biology. This work was supported by the Medical Research Council (MC_UP_1201/9). DJLDS is supported by a Medical Research Council Doctoral Training Partnership studentship (MR N013433-1). FWL is a non-stipendiary European Molecular Biology Organisation Postdoctoral Fellow (EMBO, ALTF 845-2021), supported by the Netherlands Organisation for Scientific Research (NWO-Rubicon, 019.211EN.032ß). AA is supported by a Medical Research Council Career Development Fellowship (MRC-1071140).

## Author contributions

FWL, DJLDS, AA, and LP performed experiments, analysed the data and wrote the manuscript. MAL designed and supervised the study and wrote the manuscript.

## Competing interests

MAL and LP are inventors on patents describing organoid methods, MAL is co-founder and advisory board member of a:head bio.

## Materials & methods

### Stem cell culture

Mouse E14 embryonic stem cells (ESCs; RRID: CVCL_C320) were obtained from the laboratory of Marta Shahbazi at passage 34. The line was frequently tested and confirmed as free from mycoplasma contamination. E14 ESCs were expanded in 2iLif conditions as standard, but moved toStemflex medium (Gibco #A3349401) supplemented with 1000 U ESGRO recombinant mouse leukemia inhibitory factor (mLIF; Sigma-Aldrich #ESG1107), on plates coated with growth factor reduced Matrigel (8.7 μg/cm^2^; Corning #356230), and at 37°C with 5% atmospheric CO_2_ before generation of organoids. Cells were regularly passaged every 3-4 days with 0.7 mM EDTA, or with TrypLE Express (Gibco #12604013) when individual colonies became sufficiently large to inhibit optimal growth. Human ESCs were H9 (WA09 from WiCell) and were approved for use in these studies by the UK Stem Cell Board. H9s were cultured as above, in StemFlex on Matrigel but without the addition of mLIF.

### Cerebral organoid culture

Mouse cerebral organoids were generated by following previously described unguided cerebral organoid protocols ^60^. Previously, mouse organoids were generated using a different protocol from human ^6,61^. To generate tissue comparable to human, we therefore focused on developing a method identical to the optimised method for human ^6,60,62^. First, to ensure the same pluripotent starting state, E14 ESCs were cultured for at least one passage in StemFlex as is routine for human ESCs, as described above. Beginning on day 0, E14 ESCs were dissociated to single cells with StemPro Accutase (Gibco #A1110501) and resuspended in commercial EB Formation medium (STEMdiff Cerebral Organoid kit, StemCell Technologies #8570) supplemented with 50 μM ROCK inhibitor Y27632 (SantaCruz Biotechnology #sc-281642A). 4,000 or 9,000 cells per well were seeded into ultra-low attachment U-bottom 96-well plates (Corning #7007) to form embryoid bodies. On day 2, the culture media was replaced with Induction medium (STEMdiff Cerebral Organoid kit, StemCell Technologies #8570). On day 5, embryoid bodies were embedded in droplets of Matrigel (Corning #354234), as described previously ^62^, and transferred to standard 6-well plates with Expansion medium (STEMdiff Cerebral Organoid kit, StemCell Technologies #8570). On day 6, the culture media was replaced with Improved Differentiation Medium without vitamin A (IDM-A; 1:1 [v/v] DMEM/F12 (Gibco #11330032) and Neurobasal (Gibco #21103049) media, 1:200 [v/v] N-2 supplement (Gibco #7502048), 1:50 [v/v] B-27 supplement minus vitamin A (Gibco #12587010), 1x GlutaMAX (Gibco #35050038), 0.5x MEM non-essential amino acid solution (MEM-NEAA; Gibco #M7145), 2.5 μg/mL human insulin (Sigma-Aldrich #I9278), 50 μM 2-mercaptoethanol (Gibco #31350010), 1:100 [v/v] penicillin-streptomycin (Gibco #15140122)). On day 7, Matrigel was removed from the organoids, as described below, and they were moved to 5 cm dishes on an orbital shaker (57 RPM, 25 cm orbit; 0.45 *g*) with IDM-A medium supplemented with 1:50 [v/v] Matrigel. For choroid plexus patterning, EBs were treated with 3 μM CHIR (TOCRIS, #4423) and 20 ng/ml BMP4 (Peptotech, #315-27) on day 7. For unsliced whole organoid cultures, culture media was replaced on day 8 with Improved Differentiation Medium plus vitamin A (IDM+A; IDM-A formulated with B-27 supplement (including vitamin A; Gibco #17504044) and 400 μM L-ascorbic acid (Sigma-Aldrich #A4403)) and refreshed every 4 days.

Human cerebral organoids were generated using the same methods as described above, but using the protocol timings as originally described for humans ^62,63^

### Matrigel removal

To limit the outgrowth of axons and with it the migration of cells from embedded organoids, Matrigel was depolymerised using cell recovery solution (Corning #354253) and incubation at 4 °C. First, the bulk of Matrigel was removed by dissection with sterile needles and up to 6 organoids were transferred to a 5 cm dish using a wide-bore P1000 tip. Subsequently, the media was removed, 3 mL of cold cell recovery solution was added, and the organoids were incubated at 4 °C for 20 min with regular gentle agitation to mix. Cell recovery solution was then removed and the naked organoids gently washed thrice with ice-cold phosphate-buffered saline (PBS; 125 mM NaCl, 17 mM Na_2_HPO_4_, 9 mM NaH_2_PO4, pH 7.4). Organoids were transferred to IDM-A supplemented with 1:200 [v/v] dissolved Matrigel in a 5 cm dish and incubated at 37 °C on an orbital shaker (0.45 *g*).

### Generation of air-liquid interface cerebral organoids (ALI-CO) cultures

Mouse ALI-CO cultures were generated using a similar protocol to human ALI-COs as described before ^32^. In brief, day 8-12 organoids were harvested, embedded in 3% low-gelling temperature agarose (Sigma #A9414) in molds (Sigma #E6032) and incubated for 30-60 min on ice. Sections of mouse organoids with 300 um thickness were collected using the Leica VT1000S vibrating microtome, and cultured on Millicell-CM culture inserts (Millipore #PICM0RG50) in serum-free slice culture medium (SFSCM; Neurobasal (Invitrogen #21103049), 1:50 (v/v) B27+A (Invitrogen #17504044), 1X (v/v) Glutamax (Gibco #35050038), 1:100 [v/v] penicillin-streptomycin (Gibco), 1:500 (v/v) Fungin (Invivogen #ant-fn)). After 3 days, and every 3-4 days after, ∼50% of the medium was replaced with BrainPhys medium (BrainPhys Neuronal Medium N2-A SM1 kit; StemcellTechnologies #05793), 1:100 [v/v] penicillin-streptomycin (Gibco), 1:500 (v/v) Fungin (Invivogen #ant-fn)).

### Fixation, cryosectioning and immunohistochemistry

At indicated timepoints, organoids were fixed with 4% [v/v] paraformaldehyde in phosphate buffer (145 mM Na_2_HPO_4_, 21 mM NaH_2_PO_4_, pH 7.4) for 1 h at RT for whole organoids, or overnight at 4 °C for organoid slice cultures. Organoids were then washed thrice in PBS. For cryosectioning, organoids were incubated in phosphate buffer with 30% [w/v] sucrose at 4 °C overnight. Organoids were embedded in 7.5% [w/v] gelatin (Sigma-Aldrich #G1890), 30% sucrose in phosphate buffer and the gelatin blocks frozen by submersion in cold isopentane (-50 °C) for 90 s. Frozen blocks were stored at -80 °C until being sectioned in 20 μm slices by CM1950 cryostat (Leica Biosystems) and collected on charged slides (SuperFrost Plus Adhesion; Fisher Scientific #J1800AMNZ). Cryosections were incubated with primary antibodies overnight at 4 °C, and secondary antibodies for 1 h at room temperature, with 3x 10 min PBS wash steps after every antibody incubation. Whole ALI-COs were incubated with primary antibodies and secondary antibodies for 48 h at room temperature, with 3x 8 h washes after every antibody incubation. Primary and secondary antibodies were diluted in permeabilisation buffer (0.25% Triton-X, 4% donkey serum in PBS). Primary antibodies used: N-cadherin (BD Biosciences #610920), Brachyury (R&D Systems #AF2085), E-cadherin (BD Transduction #610181), Sox17 (Abcam #ab224637), Sox10 (R&D Systems #AF2864), En2 (SantaCruz Biotechnology #sc-8111), Foxg1 (Abcam #ab18259), Otx2 (Abcam #ab21990), Pax2 (Abnova #H00005076-M01), Sox2 (Abcam #ab97959), Tuj1 (Biolegend #801213), Doublecortin (SantaCruz Biotechnology #sc-8066), Satb2 (Abcam #ab51502), Bassoon (Enzo Life Sciences, SAP7F407), Ctip2 (Abcam, #ab18465), Cux2 (Abcam, #ab216588), GFAP (Abcam, ab7260), MAP2 (Abcam, ab5392), PSD95 (Abcam, ab18258), Pan-axonal neurofilaments Clone SMI312 (Biolegend, #837904). Whole ALI-COs were mounted using Prolong Diamond (Invitrogen, P36970) with coverslips on microscope glass slides.

### EmGFP labelling

To achieve sparse labelling of cells, ALI-COs were transduced with 0.3 μl CytoTune emGFP Sendai fluorescence reporter (ThermoFisher #A16519).

### Microscopy and live-imaging

Confocal imaging of fixed samples and live-imaging was performed on an inverted Nikon CSU-W1 spinning-disk microscope, equipped with 10x/0.45 NA Air, 25x/1.05 NA Silicone Oil, 60x/1.2 NA Water lenses, sCMOS camera (95% QE), and a heated and gas-controlled incubation chamber. For live-imaging, ALI-COs were kept at 37 °C and 5% CO2 and imaged through the optically translucent Millicell-CM culture inserts. Timelapses of axon outgrowth were acquired with a 10 min interval and gentle exposure settings to avoid phototoxicity and axon tract repulsion. Displayed microscopy images represent a maximum intensity projection of a Z-stack covering the region of interest. Images of synaptic structures were acquired on an inverted Zeiss LSM 880 Airyscan equipped with an 63x/1.4 NA Oil lens, and GaAsp, spectral and Airyscan detectors. The super resolution settings of the Airyscan detectors were used to achieve high resolution images. Live-imaging of calcium dynamics was conducted on an inverted Nikon X1 Spinning Disk inverted microscope, containing a 60x/1.2NA Water lense, SCMOS cameras (95% QE), and a heated and gas-controlled incubation chamber.ALI-COs were kept at 37 °C and 5% CO2, and timelapses were acquired at a 1Hz imaging speed and single plane. Brightfield images of whole organoids for gross morphology were acquired using an Evos XL Core microscope (Thermo Fisher Scientific). Images of ALI-COs *in situ* on HD-MEAs were acquired using an iPhone X (Apple Inc.). Images were processed in Fiji or Imaris.

### MEA recordings and analysis

Extracellular electrophysiological recordings with high spatial resolution were performed on ALICOs using commercially available high-density multi-electrode arrays (HD-MEAs) containing 26400 electrodes of 17.5 mm pitch, in a 2.1 mm x 3.85 mm area (MaxWell Biosystems AG, MaxOne). Prior to recordings, HD-MEAs were incubated for one hour in 70% ethanol, before being washed three times with sterile water, and a further three times with BrainPhys Medium. ALICOs were prepared for recordings as follows: cell culture inserts where ALICOs were being cultured were gently flooded with BrainPhys Medium such that ALICO slices floated up from the inserts. ALICOs were collected in a P1000 pipette with a cut tip, or where too large for this were gently lifted using a small spatula, and transferred into HD-MEA wells filled with BrainPhys Medium. Media was completely removed from the well such that ALICOs rested on the HD-MEA surface, and were incubated at 37°C for 5 minutes without media, to facilitate attachment of the ALICO to the surface. A small volume of BrainPhys Medium (10-30ml) was then gently pipetted into the well – sufficient to keep the ALICO moist but not so much to cause the slice to float off the surface. A lid was placed on the HD-MEA well to prevent media evaporation, and HD-MEAs were allowed to reach a stable temperature and CO_2_ in the incubator before recordings were commenced. BrainPhys Medium used for transfer of ALICOs to HD-MEA, and during recordings, was that which the ALICOs were being cultured on immediately prior (rather than fresh media).

Recordings were performed in the incubator via wired connection and were operated and processed remotely using the MaxLab Live Server (MaxWell Biosystems AG, Version 22.1.4 e6d15224) via the MaxLab Live Scope graphical user interface. Three kinds of recording assays were performed (available pre-programmed in the MaxLab Live software) in sequence on each ALICO, each recording simultaneously from 1000 of the 26400 available electrodes at any one time to ensure good signal-to-noise ratio. Recording assays are outlined briefly below, summarised from the MaxLab Live Manual (MaxWell Biosystems, 2022. Release 22.2.4).

Activity Scan assays recorded from sparse electrodes 35mm apart, in 7 spatial configurations for 30 seconds each, to cover the whole area of the MEA with a spatial resolution of 816 electrodes/mm^2^.

Network Scan assays recorded from a 3x3 electrode block centred at the greatest amplitude location of each of the top Neuronal Units (preference: Firing Rates) simultaneously, as determined from the Activity Scan assay data of that same ALICO. Recordings were 300 seconds.

AxonTracking assays recorded continuously from a 3x3 electrode block centred at the Neuronal Units determined from the Activity Scan assay data, and from other electrodes across the MEA region of interest in multiple random configurations in order to capture simultaneous activity across the area. Recordings were approximately 30 minutes.

Analysis of ALICO electrophysiology data collected using HD-MEAs was performed primarily through the MaxLab Live Server (MaxWell Biosystems AG, Version 22.1.4 e6d15224) via the MaxLab Live Scope graphical user interface. For overviews of ALICO electrophysiological activity, via action potential spike-calling across the MEA from Activity Scan assays, the following thresholding parameters were used: 0.1Hz firing rate; 20mV amplitude; 200ms interspike interval. Further analyses of axon voltage changes and mapping were performed in MatLab as described in ^58^.

### Statistical analysis

Statistical details are included in corresponding figure legends. *P*LJvalues are annotated as follows: ns P > 0.05, \**P* < 0.05, \*\**P* < 0.01, \*\*\**P* < 0.001 and \*\*\*\**P* < 0.0001. Data processing and statistical analysis were conducted in Prism GraphPad (version 10.0) software.

---

**Supplementary Figure 1:**
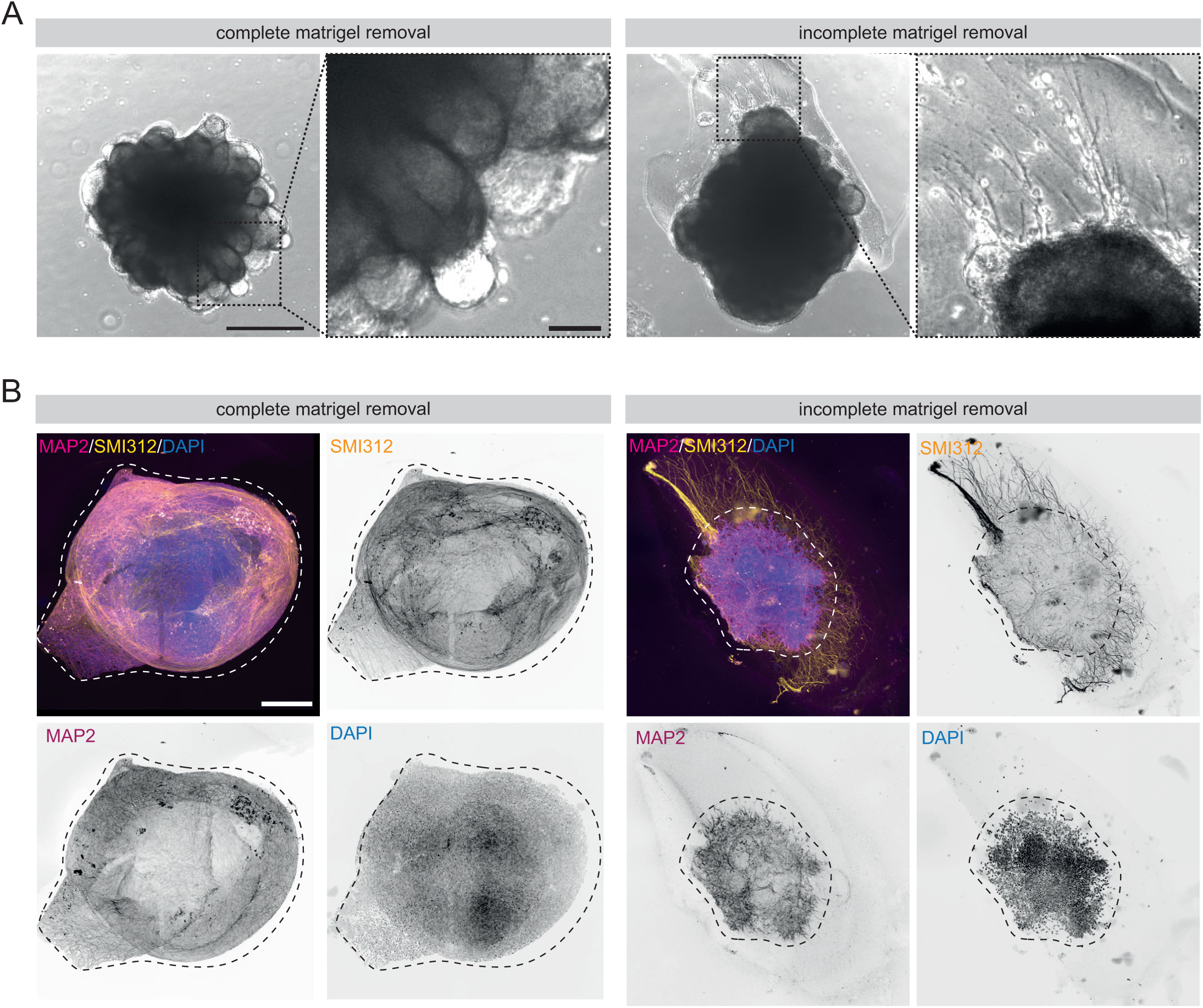
Incomplete Matrigel removal affects tissue architecture in mouse brain organoids. (A) Brightfield microscopy images of day 9 mouse brain organoids. Left: a mouse brain organoid where Matrigel was completely removed using a combination of microdissection and chemical depolymerisation, as detailed in the material and methods section. Right: a mouse brain organoid where Matrigel was only removed using microdissection, which typically results in partial removal only, leading to subsequent cell migration and outgrowth of processes into the Matrigel. Scale bars = 500 μm (full size) and 100 μm (zoom). (B) Whole-mount mouse brain organoid slice cultures immunostained for MAP2, marking neuronal somatodendritic domains, pan-axonal neurofilaments (SMI312), marking axons, and DAPI, marking nuclei. Left: a day 8+7 organoid with complete removal of Matrigel, as described in (A), with axons projecting to other regions in the organoid. Right: a mouse brain organoid at day 9+2 organoid with incomplete removal of Matrigel, as described in (A), leading to outgrowth of axons into the Matrigel instead of projecting to different regions in the brain organoid. Dotted line represents the organoid surface. Scale bar = 200 μm.

## Notes

### Summary of Updates

Author order has been updated on the bioRxiv system to match the PDF of the paper. No other changes to the manuscript have been made.

